# AtOM, an ontology model for standardizing use of brain atlases in tools, workflows, and data infrastructures

**DOI:** 10.1101/2023.01.22.525049

**Authors:** Heidi Kleven, Thomas H. Gillespie, Lyuba Zehl, Timo Dickscheid, Jan G. Bjaalie, Maryann E. Martone, Trygve B. Leergaard

## Abstract

Brain atlases are important reference resources for accurate anatomical description of neuroscience data. Open access, three-dimensional atlases serve as spatial frameworks for integrating experimental data and defining regions-of-interest in analytic workflows. However, naming conventions, parcellation criteria, area definitions, and underlying mapping methodologies differ considerably between atlases and across atlas versions. This lack of standardization impedes use of atlases in analytic tools and registration of data to different atlases. To establish a machine-readable standard for representing brain atlases, we identified four fundamental atlas elements, defined their relations, and created an ontology model. Here we present our Atlas Ontology Model (AtOM) and exemplify its use by applying it to mouse, rat, and human brain atlases. We propose minimum requirements for FAIR atlases and discuss how AtOM may facilitate atlas interoperability and data integration. AtOM provides a standardized framework for communication and use of brain atlases to create, use, and refer to specific atlas elements and versions. We argue that AtOM will accelerate analysis, sharing, and reuse of neuroscience data.

## Introduction

Brain atlases are essential anatomical reference resources that are widely used for planning experimental work, interpreting and analyzing neuroscience data^1–12^. Three-dimensional (3D) digital brain atlases^11,13–17^ are increasingly employed as frameworks for integrating, comparing, and analyzing data based on atlas-defined anatomical locations (e.g. Allen brain map, https://portal.brain-map.org/; the BRAIN Initiative Cell Census Network, https://www.biccn.org/; the EBRAINS research infrastructure, https://ebrains.eu/). These resources provide anatomical context suitable for brain-wide or region specific analysis using automated tools and workflows^18–26^ and facilitate sharing and using data in accordance with the FAIR principles^27^, stating that data should be findable, accessible, interoperable, and reusable. However, the use and incorporation of different atlas resources in such workflows and infrastructures requires that atlases, tools, and data are interoperable, with relatively seamless exchange of standardized machine-readable information.

Most brain atlases share a set of common properties, but the specifications and documentation of their parts differ considerably. Detailed versioning is not yet common practice for all atlases, and lack of specific information about changes in the terminology or anatomical parcellation make it difficult to compare atlas versions. While some gold standards have been established^28^, lack of consensus regarding the presentation, specification, and documentation of atlas contents hampers reproducible communication of locations^9^ and comparison of data that have been anatomically specified using different atlases^8,24^. Atlases and their versions need to be uniquely identifiable and interoperable to enable researchers to communicate specific and reproducible location data and integrate data across specialized neuroscience fields and modalities.

To address the lack of standardization of atlas metadata, we identified four common atlas elements, defined their relations, and created the Atlas Ontology Model (AtOM). Here we characterize the properties and relations of the elements and explain their organization in AtOM. We argue that a given set of these elements, their relations, and metadata makes up a unique version of an atlas. Furthermore, we suggest a set of minimum requirements for atlases inspired by the FAIR principles, and discuss how atlases adhering to AtOM, could accelerate neuroscience data integration.

## Results

We investigated a broad selection of mammalian brain atlases^11,13,14,16,29–37^ and identified four common elements: 1) a set of reference data, 2) a coordinate system, 3) a set of annotations and 4) a terminology. Below, we describe these atlas elements and their relations, exemplify how these elements specify unique versions of an atlas, and employ AtOM to suggest minimum requirements for FAIR brain atlases. The ontology model description is publicly available via GitHub: https://github.com/SciCrunch/NIF-Ontology/blob/atlas/ttl/atom.ttl.

### The atlas elements

The atlas elements in AtOM are the reference data, coordinate system, annotation set, and terminology (Fig. 1a-c). Each of the four elements have properties, such as identifier, species, sex, and age, specified with detailed metadata (Fig. 1d).

**Figure 1.**
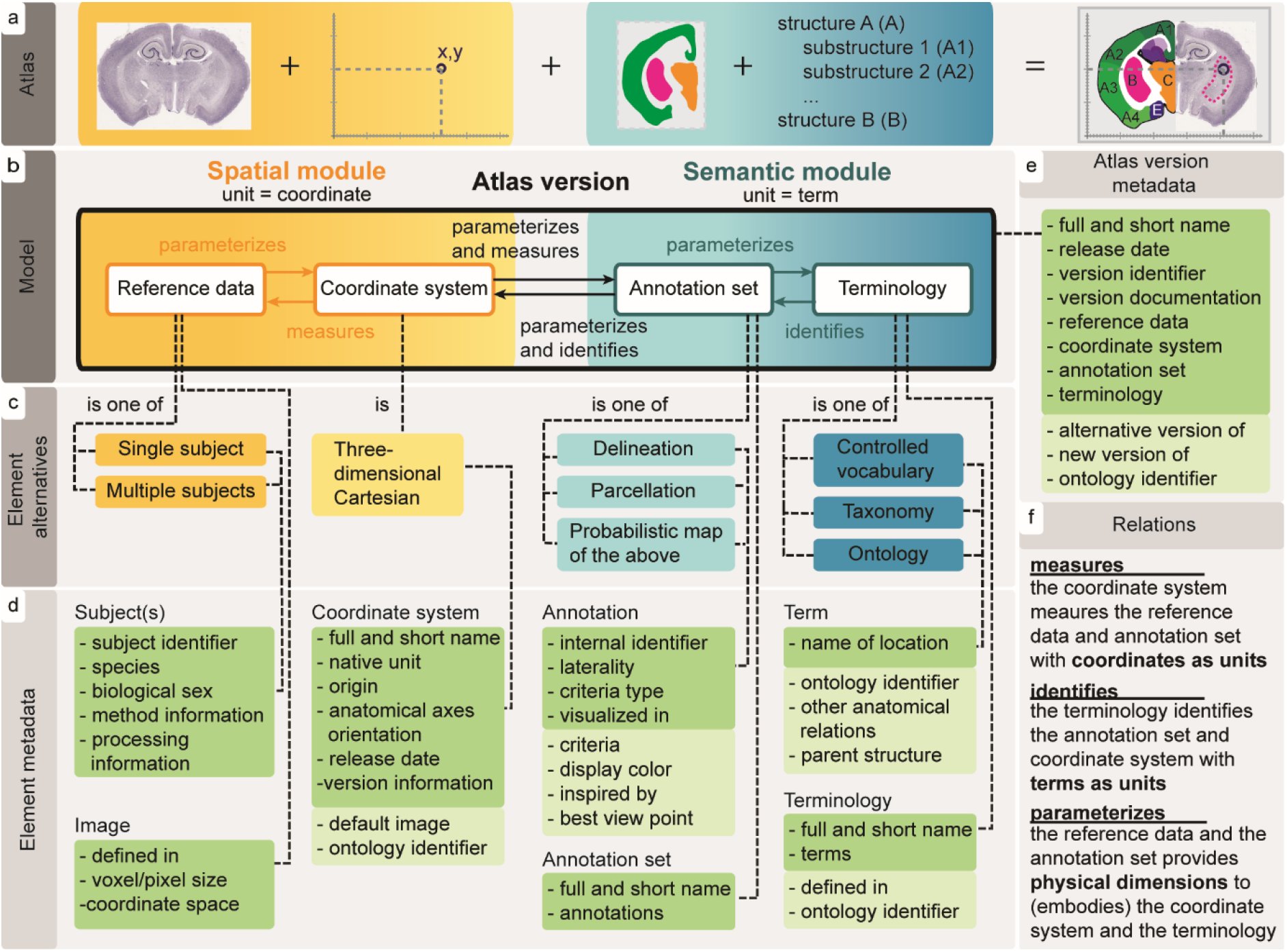
AtOM: Brain atlas elements, relations and metadata. (**a**) A diagram showing a fictional atlas divided into parts. Nissl stained coronal Platypus *(ornithorhynchus anatinus)* brain section^71^. (**b**) The Atlas Ontology Model (AtOM) showing the elements: *reference data, coordinate system, annotation set, and terminology,* and their relations (as seen in (**f**)). The model consists of two reference modules: *spatial* (containing the coordinate system and reference data, yellow) and *semantic* (containing annotations and terminology, blue). (**c**) Each element can be one of a set of alternatives, (**d**) which have a set of minimum (dark green) and additional metadata (bright green). (**e**) The aggregated atlas version metadata, and (**f**) specification of model relations; *measures* (to provide a metric to)*, identifies* (to recognize, establish or verify the identity of something) *and parameterizes* (to set the conditions of its operation).

The *reference data* of a brain atlas are graphical representations of one or several brains, or parts of brains, chosen as the biological reference for that atlas. The reference data often consist of histological or tomographic images. These images reflect different biological features of a selected specimen^14,17,32,33^, a set of different subjects representing different features and image orientations^38^, or a population average^11,13,16^. The level of detail and size of brain regions that can be identified is determined by the spatial resolution of the reference data. For example, the widely adopted human reference datasets of the Montreal Neurological Institute (MNI)^39,40^ are based on averaged magnetic resonance imaging (MRI) scans and represent suitable reference data for macroanatomy, while the single-subject *BigBrain* model^31^ provides a reference dataset for identification of cortical layers and more fine-grained cortical and subcortical structures^17^.

The *coordinate system* of an atlas provides a framework for specifying locations with units, origin, direction, and orientation. The coordinate system is usually, but not always, a 3D Cartesian coordinate system. Examples of coordinate systems which go beyond a 3D Cartesian system are spatio-temporal systems, with additional time and surface dimensions^41^. In neuroscience, many coordinate systems are defined using characteristic features of the skull^32,33^ or specific anatomical landmarks identified within the brain^14,42^.

The *annotation set* of an atlas consist of graphical marks or labels referring to spatial locations determined by features observed in, inferred from, or mapped onto the reference data, specifying structures or boundaries. An annotation set may demarcate anatomical boundaries or regions with lines, fully delineate them with closed curves^11,14,32,33^, or directly label coordinates with brain structures in the form of volumetric or surface maps. In the case of probabilistic maps, coordinates are labelled with the probabilities of a certain region or feature being present at a given location^16,43–45^. Probabilistic maps are typically aggregated from annotations identified in different individuals, encoding variation across a number of subjects^16^.

The *terminology* of an atlas is a set of terms that identifies the annotations, providing human readability and context, and allowing communication about brain locations and structural properties. In its simplest form, a terminology can be a list of unique identifiers, but is typically a set of descriptive anatomical terms following specific conventions. Atlases employ different terms, conventions, and approaches to organizing brain structures into systems based on the methodology used to create them as well as their intended use cases. For example, some use developmental organization^46,47^, while others use brain systems^37^, microstructural organization^17^, multimodal features^48^, or are specialized for particular brain regions^49,50^. An atlas terminology may be a controlled vocabulary (flat list), a taxonomy and partonomy (hierarchical list), or an ontology (hierarchy and additional axioms).

### Relations among the elements

The four elements of AtOM have specific relations (specified in Fig. 1f), sorted into a *spatial module,* consisting of the reference data and the coordinate system (Fig. 1b, yellow), and a *semantic module,* consisting of the annotation set and the terminology (Fig. 1b, blue).

The elements of the *spatial module* provide the physical and measurable dimensions of the atlas. The biological dimensions of the reference data give the conditions of operation for (i.e., *parameterize)* the coordinate system. The coordinate system provides a metric for (i.e., *measures*) the reference data, specifying the origin, orientation, and units (Fig. 1f). Coordinates are the means to derive measurements, indicate directions and spatially locate features in the reference data. The coordinate system also *measures* the annotation set, and thus connects the annotations to the features of the reference data.

The elements of the *semantic module* provide semantic identities for the atlas. The annotation set *parameterizes* the terminology in the spatial domain according to or inspired by the reference data. The terminology provides terms to establish the identity of (i.e., *identifies*) each annotation (Fig. 1f). While anatomical terms are not unique identifiers (see Atlas versioning below), they provide a means to semantically address annotations and conveying neuroanatomical knowledge and context (Fig. 1f). In this way, the terms are semantic units suitable for navigating the atlas annotations, while annotations capture the scholarly interpretations and knowledge underlying the experimental and anatomical criteria used to make them (parcellation criteria). Further, the annotation set propagates the semantic identities from the terminology, and thus semantically *identifies* locations in the coordinate system.

The relations of the atlas elements are pathways for translating information between the spatial and semantic modules. A researcher may consult an atlas to observe the physical shape and location associated with a given anatomical term, or to identify the anatomical term assigned to specific coordinates, or biological features observed in the reference data. Thus, the model is a continuous, bidirectional loop providing several starting points for researchers to translate and compare information across atlas elements.

### Atlas versioning

With an overview of the elements and relations of AtOM in hand, we are now in position to examine how they facilitate clear versioning of an atlas. In AtOM, an atlas version is a concrete instance of an atlas, and consists of specific elements, relations, and metadata (Fig. 1). Figure 2 and Table 1 shows the most recent versions of the EBRAINS supported mouse^11^, rat^14^, and human^16^ brain atlases modeled using AtOM. An important consequence of AtOM is that the atlas version changes if there are alterations to any element. Examples of alterations include revising annotations or terms, modifying the reference data or coordinate system, or replacing an element. Such changes have consequences for the specific properties and use of an atlas, and should be specified as a new atlas version. The changes made from one version to another can be described in atlas version documentation, and new versions of an atlas are usually distinguished by a new version name. The simplest way to do this is by iterative version numbering. Table 2 shows a complete overview of all versions of the Allen Mouse Brain Atlas Common Coordinate Framework (AMBA CCF)^11,13^, the Waxholm Space atlas of the Sprague Dawley rat brain (WHS rat brain atlas)^14,36,37^, and selected alternative versions of the Julich-Brain Cytoarchitectonic Atlas (Julich-Brain Atlas)^16^. In the last versions of the AMBA CCF (v3 2015-2017)^11,13,30,51–53^ and the WHS rat brain atlas (v1.01-v4)^14,29,36,37^ the semantic elements (annotation set and terminology) have been changed across versions, while the spatial elements (reference data and coordinate system, Table 2) have been kept constant. This continuation across versions allows translation of information and experimental data registered to the reference data are compatible with all versions of the mouse and rat atlas versions.

**Figure 2.**
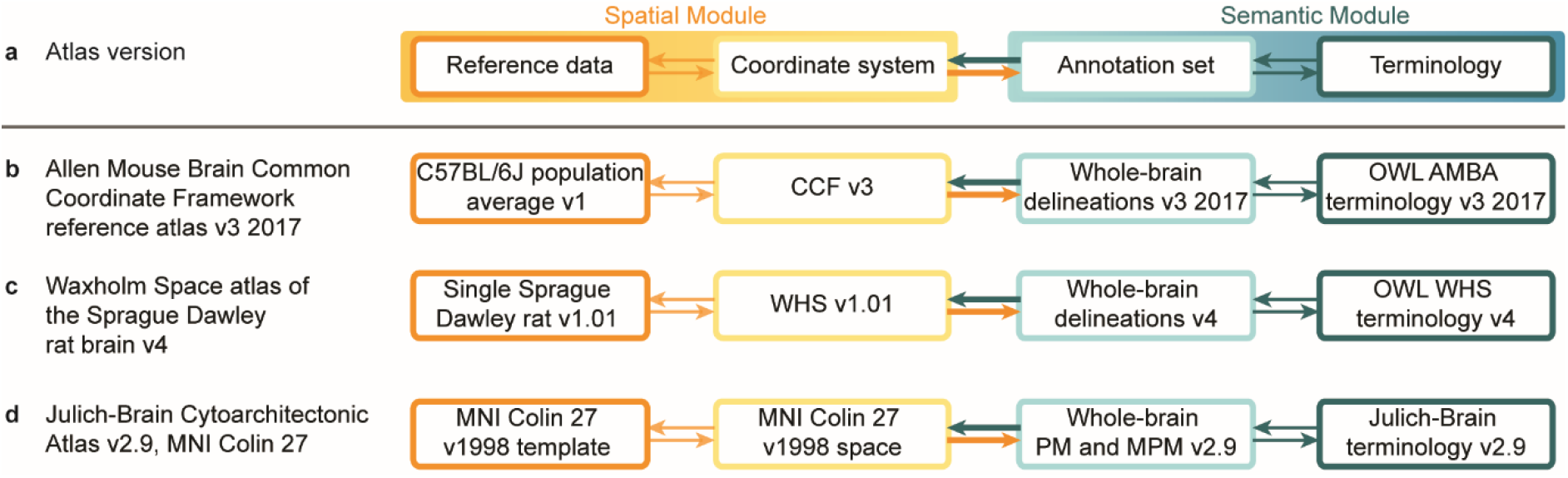
AtOM representation of the most recent EBRAINS supported mouse, rat, and human brain atlas versions. (**a**) Diagram showing AtOM. (**b-d**) Tabular view of the most recent versions of (**b**) the Allen Mouse Brain Atlas Common Coordinate Framework^11^, (**c**) the Waxholm Space atlas of the Sprague Dawley rat brain^14^ and, (**d**) one alternative representation of the Julich-Brain cytoarchitectonic atlas^16^, which are all accessible in the EBRAINS infrastructure (https://ebrains.hbp.eu/services/atlases). A more detailed representation of these atlas versions can be found in Table 1. Table 2 show all version of the mouse and rat atlases, as well as all the alternative representation of the human brain atlas v1.18 and v2.9. CCF, Common Coordinate Framework; OWL, Web Ontology Language; AMBA, Allen Mouse Brain Atlas; WHS, Waxholm Space; MNI, Montreal Neurological Institute; PM, probabilistic maps; MPM, maximum probability maps.

**Table 1.** Mouse, rat and human brain atlas version metadata.

|  |  |  |  |
| --- | --- | --- | --- |
| Full name | Allen Mouse Brain Atlas Common Coordinate Framework v3 2017 | Waxholm Space atlas of the Sprague Dawley rat brain v4 | Julich-Brain Cytoarchitectonic Atlas v2.9, MNI Colin 27 |
| Short name | AMBA CCF v3 2017 | WHS rat brain atlas v4; WHSSDv4 | Julich-Brain v2.9, Colin 27 |
| Version identifier | 3, 2017 | 4 | 2.9, Colin 27 |
| Version innovation | Publication<sup>11</sup>; White paper AMBA CCF v3 2017 (http://help.brain-map.org/display/mouseconnectivity/Documentation) | Publication<sup>14</sup>; Webpage (https://www.nitrc.org/projects/whs-sd-atlas) | Publication<sup>16</sup>; EBRAINS datasets<sup>54,55</sup> |
| Alternative version of | NA | NA | Julich-Brain v2.9, MNI 152; Julich-Brain v2.9, BigBrain; Julich-Brain v2.9, fsaverage |
| New version of | AMBA CCF v3 2016 | WHS rat brain atlas v3.01 | Julich-Brain v2.5, Colin 27 |
| Release date | NA | 01.10.2021 | 31.07.2021 |
| Reference data | C57BL/6J population average v1 | Sprague Dawley rat v1.01 | MNI Colin27 v1998 template |
| Coordinate system | CCF v3 | WHS v1.01 | MNI Colin27 v1998 space |
| Annotation set | Whole-brain parcellation, v3 2017 | Whole-brain parcellation, v4 | Whole-brain probabilistic maps and maximum probability maps |
| Terminology | OWL AMBA CCF terminology, v3 2017 | OWL WHS SD terminology, v4 | Julich-Brain terminology, v2.9 |
| License | Not available, but see legal note (https://alleninstitute.org/legal/citation-policy/) | Creative Commons Attribution-ShareAlike (CC BY-SA) 4.0 | Creative Commons Attribution-NonCommercial-ShareAlike (CC BY-NC-SA) 4.0 |

AMBA, Allen Mouse Brain Atlas; CCF, Common Coordinate Framework; MNI, Montreal Neurological Institute; OWL, Web Ontology Language; SD, Sprague Dawley; WHS, Waxholm Space.

**Table 2.**
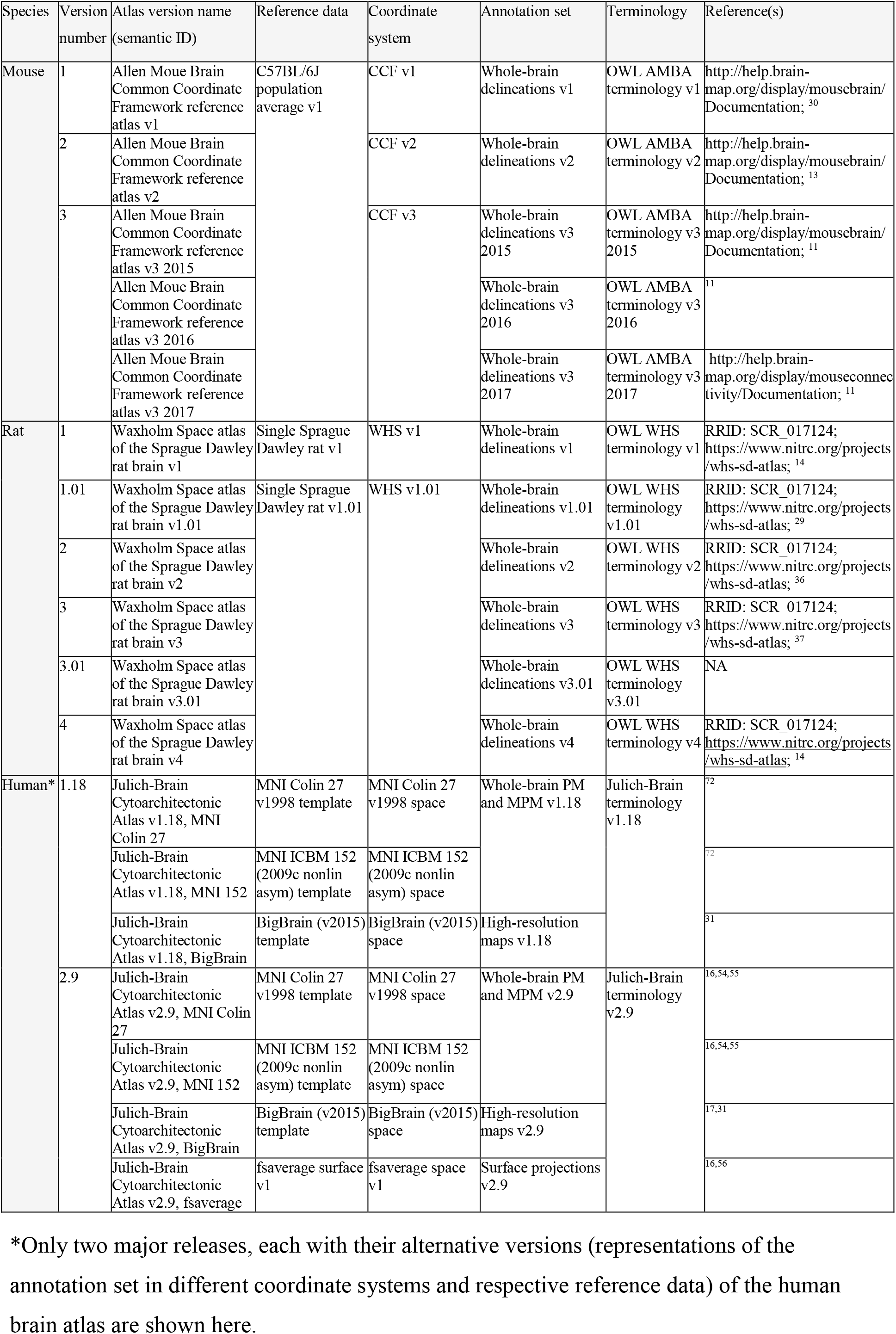
EBRAINS supported mouse, rat and human brain atlas versions.

To clearly reference a specific atlas version or AtOM element, it needs a unique identifier (ID). This is particularly important when combining different versions of elements into alternative atlas versions. The major release v2.9 of the Julich-Brain Atlas (Table 2) has four alternative versions due to its use of four complementary spatial modules: the “MNI Colin 27” (individual specimen, 1 mm resolution), “MNI 152” (population average, 1mm resolution), “BigBrain” (individual specimen, 20 μm resolution) and “fsaverage” (cortical surface representation)^17,31,54–56^. These alternative versions are identified by combining the major release identifier (v2.9) with the abbreviated name of the respective reference data and coordinate systems. Unique identifiers are also important to differentiate between identical terms, which are often similar, but not identical, anatomical areas within and across species and atlases. Ambiguity can be avoided by indexing atlas version specific terms and providing unique ontology IDs defining their properties and relations. Following AtOM, an atlas version should have unique IDs for each element and their instances, which together with version documentation facilitate clear referencing of atlas versions and specific atlas elements (Fig. 1e).

### Minimum requirements for FAIR brain atlases

Atlases are a type of research data and thus can be evaluated using the foundational principles of the FAIR guidelines^27^. These principles state that data should be findable, accessible, interoperable, and reusable through both human and machine-driven activities. Similar to experimental data, atlases can support these principles through use of unique identifiers, specific metadata, open protocols, and clear usage licenses. Furthermore, interoperability and reuse of data also requires use of “formal, accessible, shared, and broadly applicable language for knowledge representation”, as well as metadata providing detailed descriptions. Based on our proposed ontology model, we suggest the following set of four minimum requirements for FAIR brain atlases: 1) machine readable digital components, 2) defined spatial and semantic modules with element metadata, 3) specification of element versions with detailed documentation, and 4) defined element relations and metadata (Fig. 1d-e). We elaborate on these requirements below.

First, *machine-readable digital atlas components* imply that all files and metadata are available in open and non-proprietary file formats suitable for direct processing by a machine. The files and metadata for all the atlas versions shown in Figure 2 are available online, either on public websites, domain repositories, or at the atlases’ respective homepages. Table 1 shows brain atlas version metadata for the four brain atlas versions shown in Figure 2.

Second, *defined spatial and semantic modules* in an atlas mean that all elements are identifiable and accessible with clear metadata. This makes atlases easier for users to understand and easier to incorporate into tools and infrastructure. At a minimum, this can be clear naming of the essential files or documentation about the location of all necessary information (Table 1). For example, all the files needed for using the WHS rat brain atlas are available via a domain repository (Table 2).

Third, *clear versioning with granular documentation* that state all changes differentiating two version of an atlas are needed to adhere to open science and FAIR principles. Currently this is achieved through use of persistent identifiers for publications, International Standard Book Numbers (ISBN) for atlases published as books, and Digital Object Identifiers (DOI) or Research Resource Identifiers (RRID)^57^ for digital atlases. In addition, atlas reference data are made available as associated files^38^, as downloadable internet resources^11,16,17,37^, or by providing selected methodological descriptions in publications^14,16^. Some atlases also provide documentation as a list, or as text describing new features or a high-level inventory of changes. Ideally, clear versioning of an atlas should enable novice users to identify the differences between two versions (Table 2).

Fourth, the *explicit relations between atlas elements,* such as parcellation criteria and coordinate system definitions, provide an empirical foundation for translating information across the elements. This allows users to connect data to different atlas elements (semantic or spatial), and automated search or comparison of data using terms and coordinates. Traditionally, such methodological information is presented in publications^14,16^, but can also be available as white papers via a webpage^53,58,59^ or as single or distributed data publications^55^ (Table 2).

Brain atlases that fulfill these four requirements are thus expected to be sufficiently well defined to be incorporated into research infrastructures and enable automated transfer of information across atlases and between data registered to other FAIR atlases.

## Discussion

We have identified spatial and semantic elements of brain atlases, defined their relations, and created an Atlas Ontology Model (AtOM), specifying human and machine-readable metadata. Even though the AtOM elements are readily recognized in different atlases, they are often named according to traditions or common practice. For example, the reference data and the coordinate system are often considered as one entity, and referred to as the common coordinate space, reference template, reference space, brain model or atlas^7,40^. The term atlas is invariably used to address reference data, an atlas version, any of a series of atlas versions or the annotation set. The annotation set, often in combination with the terminology, has also been called parcellations, segmentations or delineations^16,17,37,43^.

Some of the AtOM elements have been suggested earlier^7^, as well as similar approaches to versioning and atlas organization^16^. However, AtOM is the first model for standardizing the common elements of any brain reference atlas, their definitions, and metadata, creating a standard to organize and share information about atlases or as a template to create an atlas.

When implemented, AtOM will facilitate precise and unique referencing of parts of an atlas, as well as the incorporation of atlases in digital tools or workflows. AtOM further provides a basis for specifying minimum requirements for brain atlases to comply with the FAIR principles. Below, we discuss how AtOM may contribute to increase interoperability among atlases, enable more standardized use of brain atlases in computational tools, and advance FAIR data sharing in neuroscience.

Interoperable atlases allow for exchange and translation of information across atlases, tools and data. Experimental data generated by different researchers typically relate to an atlas via spatial coordinates or anatomical terms, often defined by visual comparison of images or use of other observations such as measurements of functional properties. Researchers translate between the semantic and spatial location information using human readable metadata. At the same time, automated translation can be enabled via standardized, machine-readable files specifying properties and relations among atlas elements. The translation of information is dependent on interoperability across atlas elements, which can be specified at three levels: practical, technical, and scholarly.

At the *practical level,* translation of information across atlas elements is essential for interpretation and communication of anatomical locations, such as relating machine-readable coordinates to human-readable brain structure names. The relations specified between atlas elements and the defining metadata allow comparisons of annotations and terminologies across atlases representing different species or strains, developmental stages, or disease states. By aligning reference data or coordinate systems of two different atlases, information can be directly compared or translated. However, reproducible use of atlas resources depends on unambiguous citation of atlas versions. When the atlas version reference is ambiguous, or if anatomical names are given without specification of the employed atlas version terminology, it is difficult to compare location between datasets^9^. Versioning, documentation, and clear references are therefore essential for atlases that change over time.

At a *technical level*, atlas information can be accessed using computational tools, requiring specification of essential parameters and versions, such as file formats and other technical metadata. Atlases that have closed proprietary file formats may technically be digital, but without being fully machine accessible and interoperable, they are difficult to utilize in analytic tools and infrastructures.

At a *scholarly level,* anatomical parcellation and terminology should be comparable across atlases. The lack of consensus about terminologies, parcellation schemata, and boundary criteria among neuroanatomists is a major challenge for the development, use, and comparison of brain atlases^60–67^. Following different traditions, knowledge, and criteria, both domain experts and non-expert researchers may inevitably convey subjective and sometimes irreproducible information that is difficult to document. AtOM provides a foundation for organizing and communicating specific information about brain atlases in a standardized way that allows researchers to more precisely describe their interpretations, and thus contribute to increased reproducibility of results.

The value of interoperable atlases is substantial, allowing data integration, analysis and communication based on anatomical location. Brain atlases incorporated in various analytical tools open the possibility for efficient approaches to analyzing, sharing, and discovering data. For example, by analyzing images mapped to an atlas, the atlas information can be used to assign coordinates and terms to objects of interest^8,68^. Data from different publications analyzed with the same atlas are comparable, and data registered to the spatial module (reference data and coordinate system) of an atlas may also be re-analyzed with new or alternative annotation sets. Perhaps more importantly, by specifying the AtOM elements as standardized machine readable files, it becomes possible to incorporate different atlases as exchangeable modules in analytic tools and infrastructure systems^20–22,25,26^. Tools and systems using interoperable atlases can exploit the defined relations among the elements for automated operations, like data queries, calculations, or assignment of location identity to experimental data that have been associated with an atlas by spatial registration or semantic identification.

AtOM has been implemented in SANDS (spatial anchoring of neuroscience data structures, https://github.com/HumanBrainProject/openMINDS_SANDS), an openMINDS metadata model extension. The openMINDS metadata framework (https://github.com/HumanBrainProject/openMINDS, https://wiki.ebrains.eu/bin/view/Collabs/openminds/) is adopted by the EBRAINS infrastructure to describe neuroscience research products, such as data, models and software, as well as the EBRAINS atlas resources. The multilevel human brain atlas (https://ebrains.eu/service/human-brain-atlas/), an atlas framework that spans across multiple spatial scales and modalities hosted on the EBRAINS infrastructure, exemplifies how several reference data, coordinate systems, and annotation set, developed over time, can be seamlessly incorporated and presented to users through a single viewer tool. A growing repertoire of tools, services and workflows within and outside of the EBRAINS infrastructure rely on formal descriptions for automated incorporation of research products, including brain atlases and common coordinate spaces. AtOM provides a framework for keeping track of the complex relations among these resources and research products.

In conclusion, the primary value of AtOM is that it establishes a standardized framework for developers and researchers using brain atlases to create, use, and refer to specific atlas elements and versions. Atlas developers can use the model to create clearly citable and interoperable atlases. For developers incorporating atlases in tools, AtOM defines atlas elements as modules that can be seamlessly exchanged to accommodate atlases for other species or developmental stages, or to switch between versions, coordinate systems, or terminologies. By standardizing the communication and use of fundamental reference resources, we are convinced that AtOM will accelerate efficient analysis, sharing and reuse of neuroscience data.

## Methods

Ontologies are used in information sciences to specify formal representations that define the naming, properties, and relations among data and other elements that constitute a given subject or concept^69^. By specifying the relations and hierarchies of objects and processes in an ontology model, it becomes possible to create systematic and coherent links among data files, metadata, and process descriptions of relevance for a complex system. Most importantly, they enable automated retrieval of information in using computational tools^70^.

The first draft of AtOM (at the time called parcellation.ttl) was developed by eliciting requirements and use cases from the Blue Brain Project (https://github.com/SciCrunch/NIF-Ontology/issues/49). In order to ingest atlas terminologies into the NIF standard ontology a python module (https://github.com/tgbugs/pyontutils/tree/master/nifstd/nifstd_tools/parcellation) was written to convert from a variety formats into OWL. An initial version of the core ontology and 24 atlas terminologies were created. These ontologies were loaded into SciGraph (https://github.com/SciGraph/SciGraph) and queries (https://github.com/SciCrunch/sparc-curation/blob/67b534a939e2a271050c6edad97c707d8ec075d3/resources/scigraph/cypher-resources.yaml#L51-L267) were then written against the original data model using the Cypher query language in order to find atlases, terminologies, and individual terms for specific atlases, species, and developmental stages. These queries have been used in production systems for over 4 years. During this time additional atlases were ingested using the python module (now totaling 40) and an initial draft of the conceptual model for AtOM was developed (https://github.com/SciCrunch/NIF-Ontology/blob/master/docs/brain-regions.org). For a full record of the iterative development of the model to fully distinguish the major elements found in the current version (though not under their current names) see https://github.com/SciCrunch/NIF-Ontology/issues/49.

A second round of development involved further requirements collection in the context of atlas creation and the conceptual model was heavily revised, regularized, and extended in the context of the atlasing needs of the Human Brain Project (HBP) (https://github.com/SciCrunch/NIF-Ontology/commits/64c32abed9963073fab90dd5901d806fd8503da2 commit history from work during the HBP meeting in Oslo in November 21-22 2019) and the Allen Institute for Brain Sciences (https://github.com/SciCrunch/NIF-Ontology/commit/a40a8c786529f5b2e2a3a8007776d057c5830d2d, other interactions occurred, but do not have public records of their occurrence). Various iterations of the model were applied to a wide variety of atlases and atlas-like things, such as paper and digital atlases, ontologies, figures from publications, crudely drawn diagrams on table cloths, globes, geographic information systems, traditional cartographic maps, topological maps of the peripheral nervous system, and more. This was followed by collection of requirements and live ontology development carried out in the context of the HBP, which included alignment with the schemas of the openMINDS SANDS metadata model for reporting spatial metadata (https://github.com/HumanBrainProject/openMINDS_SANDS). The resulting ontological model was applied to a number of existing atlases, specifically the WHS rat brain atlas^14,36,37^, the AMBA CCF v3^11,13^, and the human Julich-Brain atlas^16,56^.

## Data availability

NA

## Code availability

NA

## Acknowledgements

The present work builds on our earlier contributions to development of brain atlases, neuroinformatics and ontologies in neuroscience with contributions from many researchers. The inspiration for developing the Atlas Ontology Model came through fruitful and valuable discussions at the INCF Workshop on Digital Brain Atlasing in Warzaw, 2019. We particularly thank Michael Hawrylycz, Alexander Woodward, Rembrandt Bakker, Ingvild E. Bjerke, Martin Øvsthus, Ulrike Schlegel, Stefan Köhnen, Xiao Gui, and Camilla Blixhavn for valuable discussions during the different stages of developing the Atlas Ontology Model. This work was funded by the European Union’s Horizon 2020 Framework Programme for Research and Innovation under the Specific Grant Agreement No. 785907 (Human Brain Project SGA2) and Specific Grant Agreement No. 945539 (Human Brain Project SGA3), The Research Council of Norway under Grant Agreement No. 269774 (INCF Norwegian Node, to JGB and TBL), and the Helmholtz Association’s Initiative and Networking Fund through the Helmholtz International BigBrain Analytics and Learning Laboratory (HIBALL) under the Helmholtz International Lab grant agreement InterLabs-0015 (to TD).

## Author contributions

**HK** contributed to conceiving the study, establishing and validating the model, writing the paper, and creating figures. **THG** contributed to conceiving the study, establishing and validating the model, creating and maintaining the ontology, writing the paper, and creating figures. **LZ** contributed to establishing and validating the model, and writing the paper. **TD** contributed to establishing and validating the model, and writing the paper. **JGB** contributed to establishing and validating the model, and writing the paper. **MEM** contributed to conceiving the study, establishing and validating the model, writing the paper, and supervising the study. **TBL** contributed to conceiving the study, establishing and validating the model, writing the paper, and supervising the study.

